# Functional Comparison Between Endogenous and Synthetic Notch Systems

**DOI:** 10.1101/2022.04.24.489295

**Authors:** Bassma Khamaisi, Vincent C. Luca, Stephen C. Blacklow, David Sprinzak

## Abstract

The Notch pathway converts receptor-ligand interactions at the cell surface into a transcriptional response in the receiver cell. In recent years, synthetic Notch systems (SynNotch) that respond to different inputs and transduce different transcriptional responses have been engineered. One class of synNotch systems uses antibody-antigen interactions at the cell surface to induce the proteolytic cleavage cascade of the endogenous Notch autoregulatory core and the consequent release of a synNotch intracellular domain (ICD), converting surface antigen detection into a cellular response. However, whereas the activation of endogenous Notch requires ubiquitylation and subsequent endocytosis of the ligand ICD, these synNotch systems do not seem to have such a requirement because the synNotch ligands completely lack an ICD. This observation raises questions about existing models for the Notch activation mechanism. Here, we test how different structural and biochemical factors affect the dependence of endogenous and synthetic Notch activation on ligand ICD. We compare the behavior of antibody-antigen synNotch (aa-synNotch) to that of endogenous Notch, and to a synNotch system that uses induced FKBP-FRB dimerization with rapamycin (ff-synNotch), which still requires a ligand ICD. We found that differences in receptor-ligand affinity, in the identity of the transmembrane domain, or in the presence or absence of extracellular EGF repeats cannot explain the differences in ligand ICD requirement that distinguishes aa-synNotch from endogenous Notch or ff-synNotch. These findings suggest that the aa-SynNotch systems bypass the ligand ICD requirement because antigen-antibody pairs are able to promote other adhesive cell-cell interactions that provide the mechanical tension needed for ligand activation.

## Scientific background

Notch signaling is a highly conserved signaling pathway promoting cellular communication between neighboring cells across metazoans. Aberrant Notch signaling is associated with various human diseases, including cancer (Aster, Pear and Blacklow, 2008), developmental abnormalities, and other pathologic conditions (Louvi and Artavanis-Tsakonas, 2012).

In mammals, there are four Notch homologues (Notch1-4) and five Delta/Serrate/Lag-2 (DSL) ligands, three of the delta-like family (Dll1, Dll3, and Dll4) and two of the jagged family (Jag1 and Jag2); Different receptors and ligands are used to activate distinct target programs, and thus both define and operate within different biological contexts (Cheng *et al*., 2007; Preuße *et al*., 2015; Nandagopal *et al*., 2018).

Notch signaling is transduced when a Notch receptor on a receiver cell binds a DSL ligand on a neighboring cell. This interaction leads to two cleavage events of the Notch receptor, one in the Notch extracellular domain (NECD) and the second in the Notch transmembrane domain (S2 and S3 cleavage events, respectively). These successive cleavage events release the Notch intracellular domain (ICD) from the membrane, allowing it to translocate to the nucleus and serve as a co-transcription factor. In parallel to the release of the Notch ICD in the receiver cell, the rest of the Notch receptor, the NECD, remains bound to the ligand and can enter the sender cell in a process called trans-endocytosis.

Notch activation requires a pulling force applied by the DSL ligands on the sender cell to the Notch receptors on the receiver cell, leading to a conformational change in the Notch regulatory region (NRR) enabling S2 cleavage (Gordon *et al*., 2015). It has been proposed that clathrin-mediated endocytosis (CME) in the sender cell is responsible for delivering this force to the ligand-receptor complex (Weinmaster and Fischer, 2011; Meloty-Kapella *et al*., 2012; Sprinzak and Blacklow, 2021).

Endocytosis of Notch ligands relies on the ubiquitylation of multiple lysine residues in the ligand ICD. In mammals, this ubiquitylation is mediated by the E3 ubiquitin ligase, Mind bomb 1 (Mib1). Some work suggests that ligand ubiquitylation recruits the endocytic adapter protein, Epsin (Langridge and Struhl, 2017), which in turn recruits the Clathrin-mediated endocytosis (CME) machinery, yet alternative lines of evidence show that DSL ligands can activate without being ubiquitylated in some cases (Berndt *et al*., 2017), and residual Notch signaling activity can be induced by Delta in the absence of Mib1 in the Drosophila wing margin (Berndt *et al*., 2017).

Recently, the core machinery of the Notch receptor has been used to develop synthetic Notch (synNotch) receptors that can receive different signals (e.g., bind specific membrane proteins in the sender cell) and transduce ectopic or customized transcriptional responses. In these systems, the natural ligand-receptor pair is replaced with the rapamycin-inducible FKBP/FRB heterodimer (Gordon *et al*., 2015) or an antibody-antigen interaction (Morsut *et al*., 2016), and the Notch ICD is replaced with a synthetic transcription factor, but the core autoregulatory machinery that allows cleavage in response to ligand binding was retained. Importantly, this synNotch has important implications in the development of cell therapy applications (Choe *et al*., 2021; Han *et al*., 2021; Tahmasebi *et al*., 2021).

Interestingly, synNotch is still active when membrane bound antigen molecules are used as synthetic ligands (aa-synNotch) even when they do not retain any intracellular sequences (aa-synNotch ligands typically have a short 8 aa ICD). While endogenous Notch requires the ligand ICD for its activation, this requirement does not seem to hold for the aa-synNotch system. Here, we sought to deduce the basis for this difference in activity by systematically evaluating the differences between the endogenous Notch system, the antibody-antigen based synNotch (aa-synNotch), and the synNotch system based on the rapamycin-induced dimerization of FKBP and FRB (ff-synNotch). We found that the aa-synNotch does not require a ligand ICD to function, whereas endogenous and ff-synNotch do. We show that the differences between the systems do not arise due to differences in receptor-ligand affinity, nor due to the specific transmembrane segment present. We also show that the presence or absence of EGF repeats in the ligand or the receptor do not explain the observed differences in behaviors between the different systems. These findings suggest that the aa-SynNotch systems bypass the ligand ICD requirement because of a differential ability of antigen-antibody pairs to promote other adhesive cell-cell interactions (e.g., clustering, membrane mobilization, etc.) that remove the requirement for ligand ubiquitylation and provide the mechanical tension needed for ligand activation.

## Results

### Endogenous ligands lacking all lysines or the entire ICD are unable to activate Notch

A number of studies have shown that Notch ligands lacking their ICD or all lysine residues in their ICD are deficient in the ability to activate Notch (Shimizu *et al*., 2002; Nichols *et al*., 2007; Heuss *et al*., 2008), even though they are present on the cell membrane and are going through endocytic processes (Daskalaki *et al*., 2011). To better understand the role of the ligand ICD, we analyzed the activity of mutant human Dll1 (hDll1) and human Dll4 (hDll4) ligands lacking all their lysine residues in the ICD or their ICD entirely. To test ligand activity, we performed a luciferase activity assay in which a Chinese hamster ovary (CHO) reporter cell line stably expressing Notch1 is transfected with a luciferase reporter, and is then co-cultured with CHO-TetR cells stably expressing different ligand variants (Figure 1A). We considered six ligands in our assays: wild type hDll1/4 (this refers to two versions, one with hDll1 and the other with hDll4), hDll1/4 lacking an ICD (hDll1/4-ΔICD), and hDll1/4 where all lysine residues were replaced by arginine in the ICD (hDll1/4-no lysine). All ligands were fused to mCherry fluorescent proteins at their C-terminus to enable visualization (all variants tested exhibited similar expression levels; Figure S1A). In line with previous observations (Nichols *et al*., 2007), we found that mutant ligands that either completely lack their ICD or have no lysines in their ICD exhibit significantly reduced Notch activation (Figure 1B).

**Figure 1.**
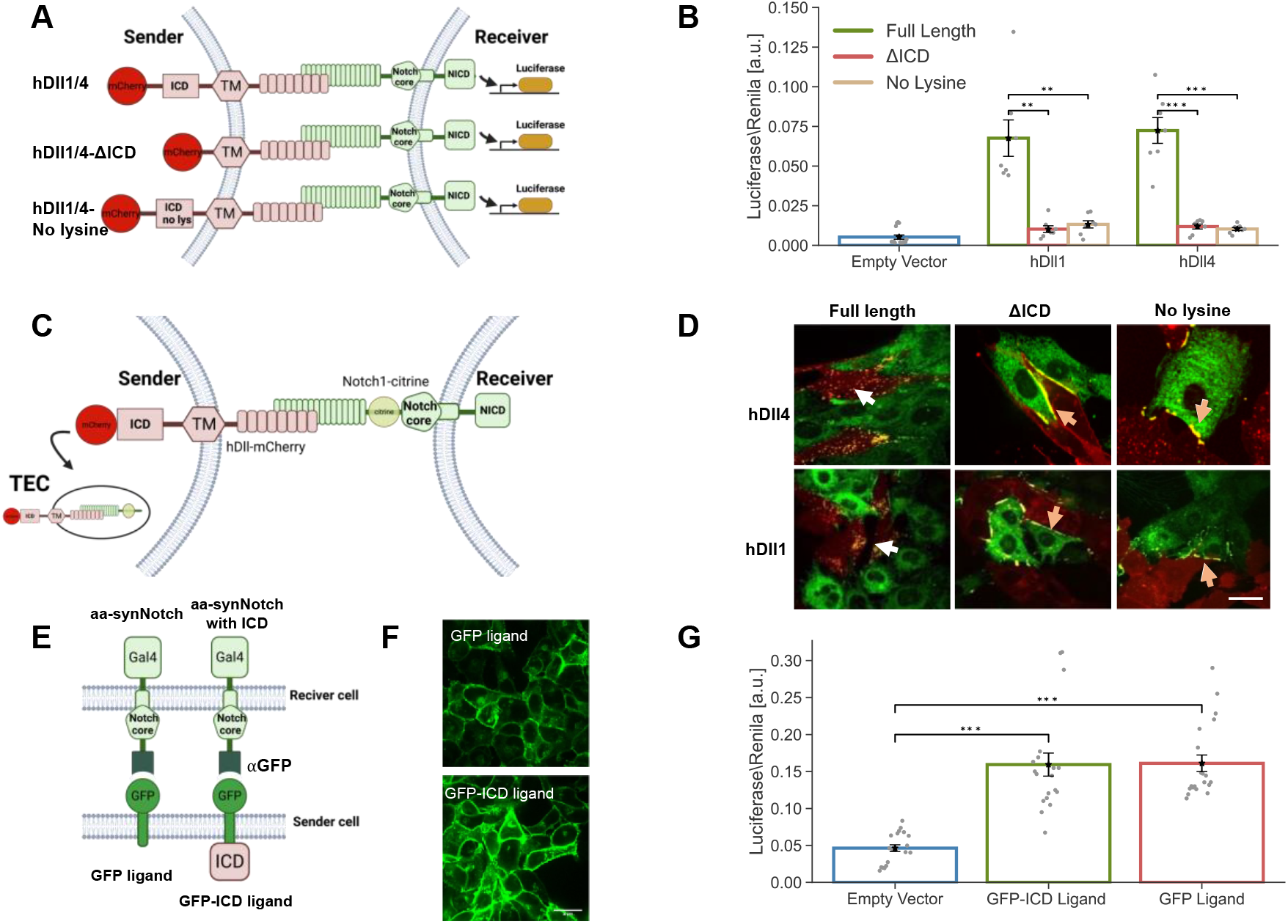
The ICD of Notch ligands is required for Notch activity in the endogenous system, but not in the aa-synNotch system. **(A)** Schematic of the Notch luciferase activity assay. In this assay, sender cells that express the Notch ligands Dll1 or Dll4 (hDll1/4) activate a Notch luciferase reporter in the receiver cells. **(B)** Luciferase activity assay showing the activation of a Notch reporter cell line (CHO-Notch1 transfected with a 12xCSL-Luciferase reporter) co-cultured with CHO-TetR cells expressing either wildtype hDll1/4 (hDll1/4), hDll1/4 lacking its ICD (hDll1/4-ΔICD), or hDll1/4 with no lysine residues in its ICD (hDll1/4-no lysine). **(C)** A schematic of the live-cell trans-endocytosis (TEC) assay where inducible hDll1/4 (and its variants) are co-cultured with Notch1 fused to citrine in its NECD (Notch1-Citrine). **(D)** Images showing a co-culture of inducible hDll1/4 variants (red) with Notch1-citrine cells (green) 10 hours after induction of ligand expression with 100 ng/ml dox. While the full-length hDll1/4 exhibits strong TEC (white arrows), both hDll1/4-ΔICD and hDll1/4-no lysine show accumulation on the boundaries (orange arrows). **(E)** Schematic of an aa-synNotch system with a receiver cell expressing an αGFP receptor, and a sender cell expressing either a GFP ligand, or a GFP ligand with hDll4ICD (GFP-ICD). **(F)** Images of U2OS cells expressing either GFP ligand or GFP-ICD. **(G)** Luciferase activity assay with U2OS cells expressing αGFP receptors co-cultured with U2OS cells expressing either GFP or GFP-ICD ligands. Data points show mean values from n=7 for (B), and n=20 for (G), from 3 and 5 independent experiments, respectively. Error bars represent S.E.M.**p<0.01, ***p<0.001. Scale bars-10μm.

To further elucidate how the tailless ligands differ from the full-length ligands at the cell surface, we also performed a trans-endocytosis assay (TEC) (Shaya *et al*., 2017). In this assay, we followed the interaction between Notch receptors and ligands by co-culturing CHO-TetR cells expressing a Notch1 receptor with a citrine tag at its ECD (Notch1-citrine) with CHO-TetR cells expressing different variants of hDll1/4. Productive signaling in this assay was manifested by the trans-endocytosis of tagged NECD into the signal sending cells (Figure 1C). Consistent with our luciferase activity assay results, these studies showed that only full-length ligands exhibited trans-endocytosis (white arrows in Figure 1D), while mutants lacking all lysine residues in their ICD or their entire ICD accumulate at the boundary of the Notch1-citrine cell but do not undergo TEC (orange arrows in Figure 1D). This observation indicates that Notch ligands lacking their lysine residues, or their ICD interact with Notch receptors but cannot activate them.

### Ligand activity does not require ligand tail in GFP-αGFP synNotch system

We next wanted to test whether signaling in an aa-synNotch system required ligand ICD. Previous work (Morsut *et al*., 2016) has shown that aa-synNotch does not require a ligand tail to be active, but did not assess whether adding a ligand tail affects signaling. To test this, we generated Human Bone Osteosarcoma Epithelial (U2OS) cells expressing either a GFP ligand (identical to the one used by Morsut *et al*., 2016) or a GFP ligand with a hDll4 tail added to its C-terminus (GFP-ICD). Sender cells expressing these ligands were co-cultured with U2OS receiver cells expressing αGFP-Gal4 synNotch receptors (single-chain GFP nanobody fused to Notch NRR and intracellular Gal4) transfected with an UAS-luciferase reporter (Figure 1E). Expression levels and membrane localization of both GFP ligands were similar (Figure 1F, Figure S1B). Measurements of luciferase activity in the receiver cells showed that the GFP ligand with and without hDll4 ICD were similarly active (Figure 1G). Thus, in contrast to the endogenous Notch system, signaling with the aa-synNotch system is not detectably affected by the presence or absence of the ligand tail.

### Receptor-ligand affinity affects the strength of activation but does not compensate for the lack of ICD

To uncover the origin of the functional differences between synthetic and endogenous systems, we systematically analyzed the molecular differences between the two systems. We first assessed whether the observed difference in the dependence on the ligand ICD stems from differences in receptor-ligand binding affinity. The binding of Notch1 to rat Dll4 has a K_D_ value of 12.7 μM for wildtype Dll4 (Luca *et al*., 2015), whereas the K_D_ value for the GFP-αGFP pair in the synNotch system is ∼50 nM (Toda *et al*., 2020), suggesting that a higher affinity might lead to a stronger ligand activity that compensates for the lack of an ICD. To test this hypothesis, we first checked whether higher ligand affinity can compensate for the lack of activity in ligands that lack their ICD. More specifically, we used several rat Dll4 (rDll4) variants that were developed recently using a yeast display library and in-vitro evolution to determine the structure of a receptor-ligand complex (Luca *et al*., 2015). These included the following variants: (i) wildtype rat Dll4 (WT) with K_D_ = 12.7 μM, (ii) the Dll4 SLP variant with K_D_ = 440nM corresponding to a 30-fold increase in affinity relative to WT Dll4, and (iii) the Dll4 E12 variant with K_D_ = 56 nM corresponding to a 225-fold increase in affinity relative to WT Dll4, an affinity that is comparable to that of the GFP-αGFP pair. We used the higher affinity ECD domains to build full-length Dll4 variants with the ICD of hDll4, and used hDll4 ECD as a reference. All ligands were fused to mCherry fluorescent proteins at their C-terminus (Figure 2A). We generated CHO-TetR stable cell lines expressing all 3 chimeric Dll4 variants as full-length proteins and in versions lacking their ICD. All variants tested exhibited similar expression levels (Figure S2), and activation by the WT chimeric ligand (rDll4ECD_WT_- hDll4ICD) was indistinguishable from the full-length hDll4 ligand. In luciferase assays, higher affinity ligands with intact ligand tails exhibited higher activity, but ligands lacking an ICD showed greatly reduced activity when compared to full-length ligands (Figure 2B).

**Figure 2.**
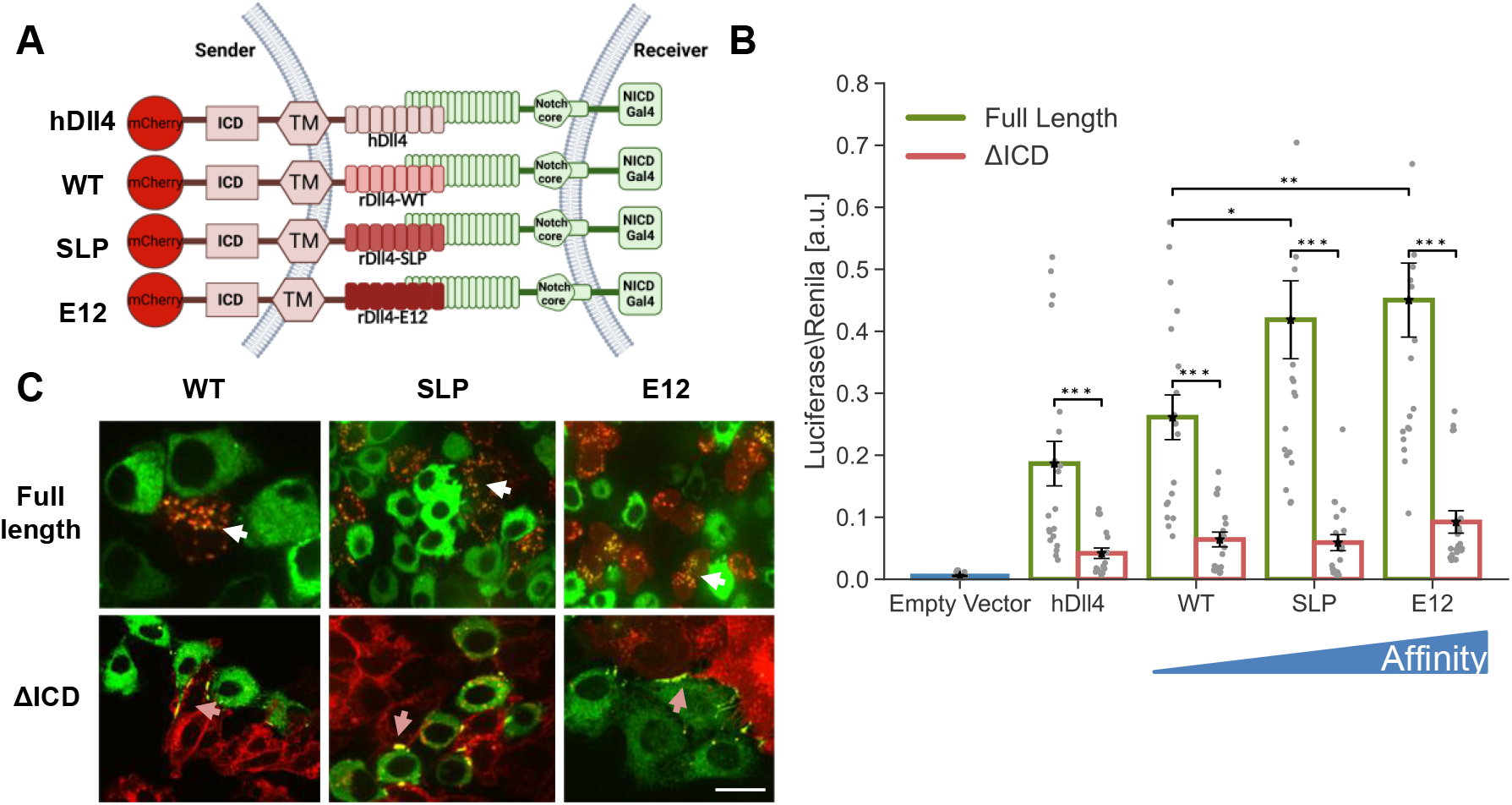
Higher affinity receptor-ligand interactions do not compensate for the lack of ligand ICD. **(A)** Schematic of the chimeric variants of Dll4 generated for testing the role of receptor-ligand affinity. Here, hDll4 is the full-length human Dll4 ligand. All the chimeric variants contain human Dll4ICD and transmembrane domains and different versions of rat Dll4ECD. The binding affinities of WT, SLP, and E12 to Notch1 are 12.7μM, 440nM, and 56nM, respectively (Luca *et al*., 2015). All the variants are placed under an inducible promoter and are fused to mCherry. **(B)** Luciferase activity assay showing the activation of a Notch reporter cell line (U2OS-Notch1-Gal4 transfected with a UAS-Luciferase reporter) co-cultured with CHO-TetR cells expressing either the indicated variants either with or without ligand ICD. **(C)** Images showing a co-culture of inducible affinity variants (red) with Notch1-citrine cells (green) 10 hours after induction of ligand expression with 100 ng/ml dox. TEC is observed in full-length ligands (white arrows). Accumulation on the boundaries, but no TEC are observed in ligands lacking their ICD (orange arrows). Data points show mean values from n=20 for (B) from 5 independent experiments. Error bars represent S.E.M. *p<0.05,**p<0.01, ***p<0.001. Scale bars-10μm.

We also tested ligand activity using the TEC assay. As with hDll4, we observed TEC with the chimeric rat full-length ligands (white arrows in Figure 2C), and only accumulation of bound Notch1 at the boundary between cells in co-cultures with ligands lacking the ICD (orange arrows in Figure 2C). Altogether, these results show that higher affinity Dll4 interactions do not compensate for the lack of ligand ICD.

### Ligand tail is required for activation of the FKBP-FRB SynNotch system

Since the GFP-αGFP affinity in the synNotch system is comparable to the affinity between Notch1 and the E12 variant (i.e., higher affinity Dll4 generated by Luca *et al*., 2015). We considered whether other specific features of the interaction domains underlie the observed differences. We therefore tested the requirement for ICD in the ff-SynNotch configuration (Gordon *et al*., 2015). This system uses the FRB domain of mTOR and the FK506 binding protein (FKBP), which interact to form a stable complex only in the presence of rapamycin. The FKBP domain replaces the Notch-binding MNNL and DSL domains (the N-terminal domains of the ligands) but retains the rest of the extracellular, transmembrane (TM), and ICD of the original Dll4. We also fused mCherry fluorescent protein at the C-terminal end of the ligands as with the endogenous ligand experiments. On the receptor, the FRB domain replaces the first 23 EGF-like repeats of Notch1, but retains the original Notch core and an ICD in which the Gal4 DNA-binding domain is substituted in place of the ankyrin repeat domain of the receptor (Figure 3A). The FKBP-rapamycin complex binds to FRB tightly with K_D_ = ∼12 nM (Banaszynski, Liu and Wandless, 2005).

**Figure 3.**
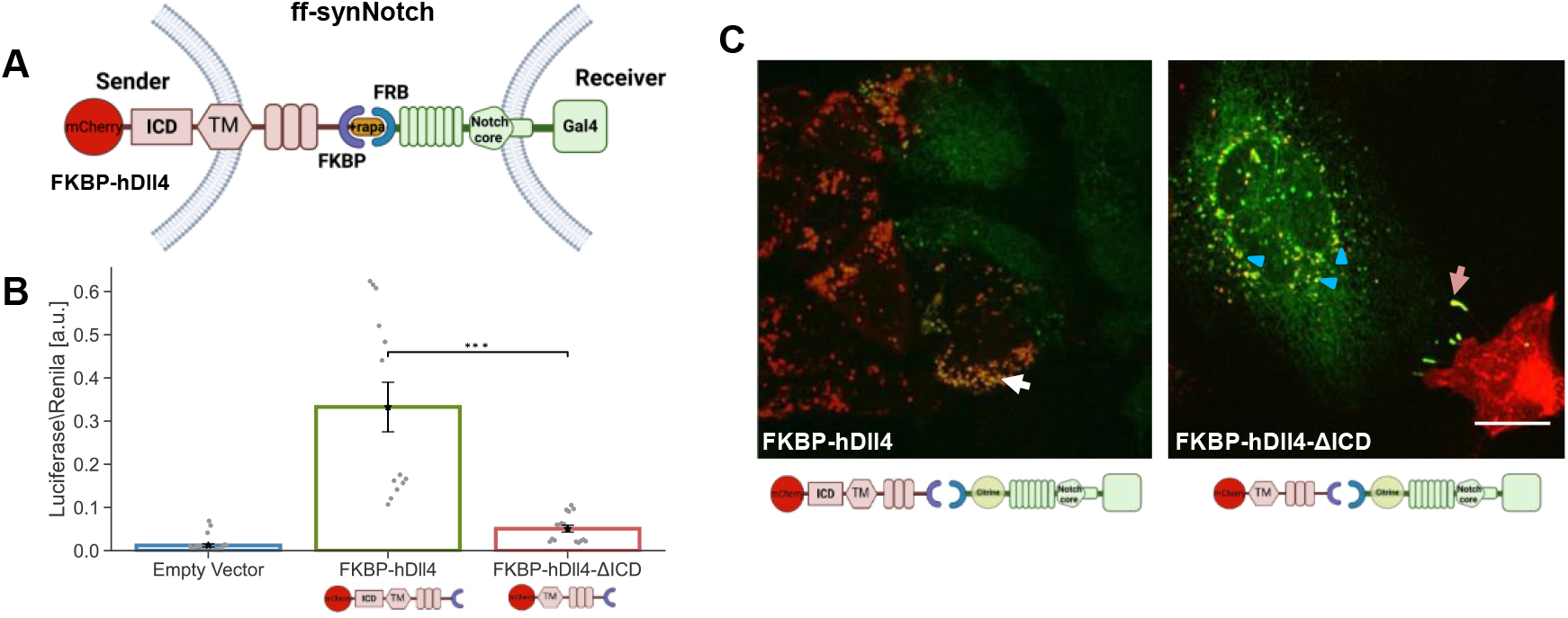
Ligand tail is required for activation of the FKBP-FRB SynNotch (ff-synNotch) system. **(A)** Schematic of the ff-synNotch system. Here, FKBP and FRB replace the N-terminal portion of Dll4 and the Notch1 EGF-like repeats 1–23, respectively (Gordon *et al*., 2015). **(B)** Luciferase activity assay showing the activation of a Notch reporter cell line (U2OS-FRB-Notch1-Gal4 transfected with a UAS-Luciferase reporter) co-cultured with CHO-TetR cells expressing the indicated variants of FKBP-hDll4 ligand either with or without ligand ICD (FKBP-hDll4-ΔICD). **(C)** Images showing a co-culture of inducible CHO-TetR cells expressing the FKBP-hDll4 or FKBP-hDll4-ΔICD (red), with U2OS cells expressing the FRB-Notch-citrine (Citrine tag is inserted in the ECD, green). Images were taken 10 hours after induction of ligand expression with 100 ng/ml dox followed by 1 hour induction of binding with 250 nM rapamycin. TEC is observed in full-length ligands (white arrows). Accumulation on the boundaries, but no TEC are observed in ligands lacking their ICD (orange arrows). Experiments with ligands lacking the ICD show reverse TEC (blue triangle). Data points show mean values from n=13 from 4 independent experiments. Error bars represent S.E.M. ***p<0.001. Scale bars-10μm.

To test activity in the ff-synNotch system, we co-cultured U2OS cells stably expressing FRB-Notch1-Gal4 with CHO-TetR sender cell lines stably expressing either full-length FKBP-ligands, or FKBP-ligands lacking their ICD. We note that variants tested exhibited similar expression levels (Figure S3). Consistent with previous work using this system (Gordon *et al*., 2015), we found that ligands lacking the ICD in this ff-synNotch system exhibited greatly reduced activity compared to the full-length ligands (Figure 3B).

We also performed a TEC assay with the ff-synNotch by inserting a citrine fluorescent tag into the ECD of the receptor (Figure 3C). As with the endogenous Notch system, we observed TEC with the ligands containing the ICD (white arrows in Figure 3C), but not with the ligands that lack the ICD. Ligands lacking the ICD showed accumulation at the cell contact boundary (orange arrows in Figure 3C) as well as reverse trans-endocytosis, in which the ligand undergoes endocytosis into the sender cell (marked by a blue triangle in Figure 3C). Such reverse TEC has been previously observed with non-functional synNotch proteins in another synNotch system in Drosophila (Langridge and Struhl, 2017). Overall, our results show that in contrast to the aa-synNotch system, the ligand ICD is required for activation in the ff-synNotch system despite both having comparable receptor-ligand affinities.

### The type of transmembrane (TM) domain does not affect dependence on ligand ICD

Since the TM region can act as an endocytosis signal in mammalian cells (González Montoro, Bigliani and Valdez Taubas, 2017), we next examined whether the different behavior of the aa-synNotch and the ff-synNotch systems can be attributed to differences in the TM domain. The TM region used for the aa-synNotch system is from the PDGFR (platelet-derived growth factor receptor), while the TM region used in the ff-synNotch system was the endogenous one from hDll4. To test whether the different behavior of the two systems depends on the TM domain, we replaced the TM of the endogenous hDll4 and FKBP ligands with the PDGFR TM and compared activities with and without the ligand ICD (Figure 4A). All variants tested exhibited similar expression levels (Figure S3). Our results, both in the luciferase and TEC assays (Figures. 4B and 4C, respectively), showed that only the full-length ligand can activate Notch receptors irrespective of the TM domain used. These findings show that the TM domain is not the source of the different behaviors of the two synNotch systems.

**Figure 4.**
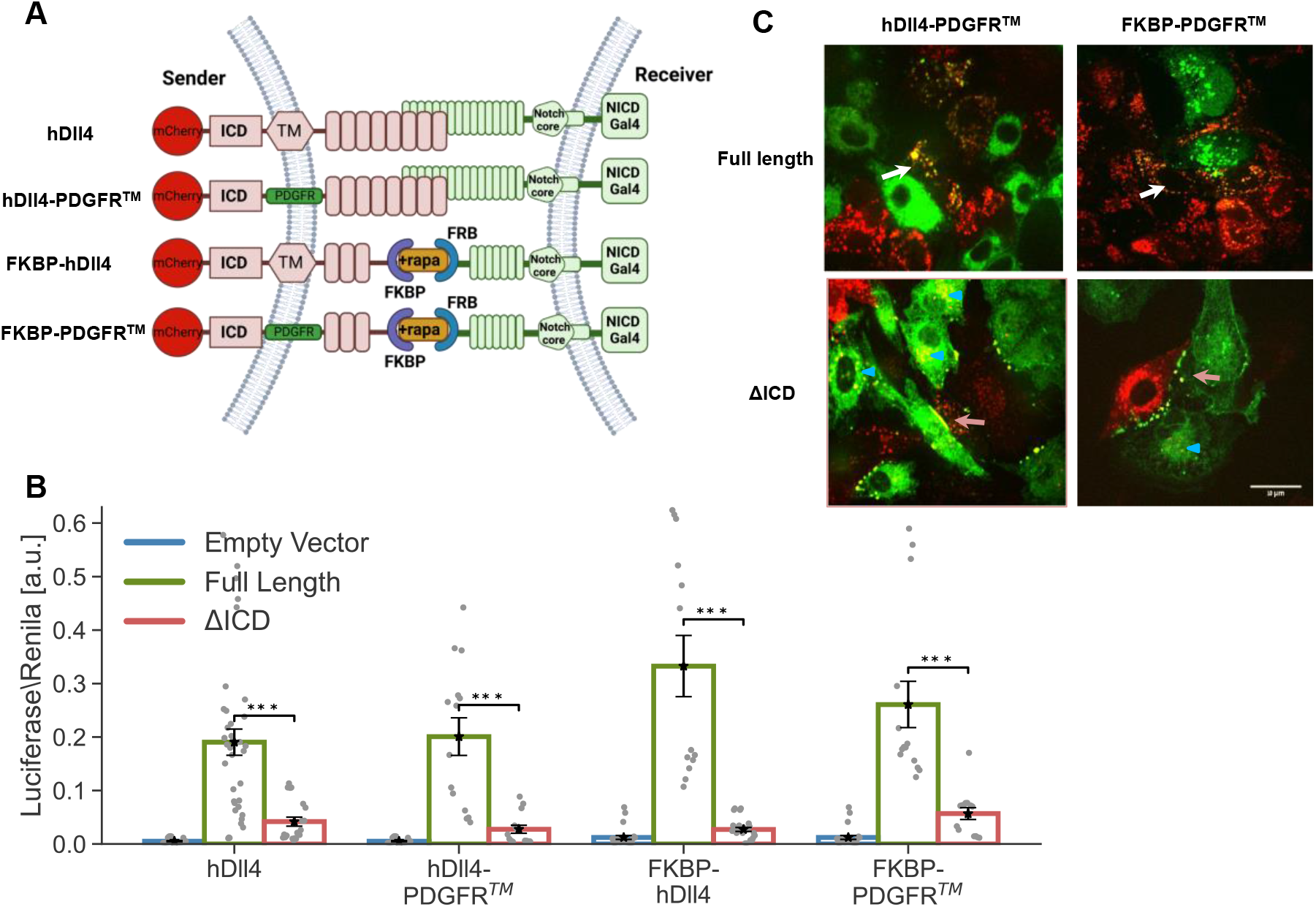
The type of TM domain does not affect dependence on ligand ICD. **(A)** Schematic of the constructs used for testing the effect of TM region on Notch activity. **(B)** Luciferase activity assay showing the activation of a Notch reporter cell line (U2OS-Notch1-Gal4 or U2OS-FRB-Notch1-Gal4 transfected with a UAS-Luciferase reporter) co-cultured with CHO-TetR cells expressing the indicated variants either with or without ligand ICD. **(C)** Images showing a co-culture of inducible CHO-TetR cells expressing the ligand variants (red) with U2OS cells expressing the Notch1-Citrine or FRB-Notch1-Citrine (green). Images were taken 10 hours induction of ligand expression with 100 ng/ml dox followed by 1 hour induction of binding with 250 nM rapamycin (only for the ff-synNotch system). TEC was observed with full-length ligands (white arrows). Accumulation at the boundaries of cell contact without TEC was observed with ligands lacking their ICD (orange arrows). Experiments with ligands lacking the ICD show reverse TEC (blue triangle). Data points show mean values from n=14 from 4 independent experiments. Error bars represent S.E.M. ***p<0.001. Scale bars-10μm.

### Removing the EGF repeats from the ff-synNotch ligands and receptors does not compensate for the lack of ICD

Another difference between the two synNotch systems resides in the extra EGF repeats in both the ligands and receptors of the ff-synNotch system. Since the distance between the receptor binding epitope and the membrane has been shown to modulate receptor-ligand interactions in other systems (James *et al*., 2008, p. 22), we reasoned that these extra EGF repeats could lead to an increase in the distance between the sender and receiver cells and thus modulate synNotch receptor-ligand interactions. To test the role of the ligand EGF repeats, we created four ligand FKBP variants with the PDGFR TM region by either including or excluding EGF repeats in the ligand ECD in full-length or tailless-ligand versions (Figure 5A). The variants tested exhibited similar expression levels (Figure S3). Both luciferase and TEC assays with these ligands show that removing the EGF repeats from the ligands did not compensate for the lack of the ligand ICD (Figures 5B-C).

**Figure 5.**
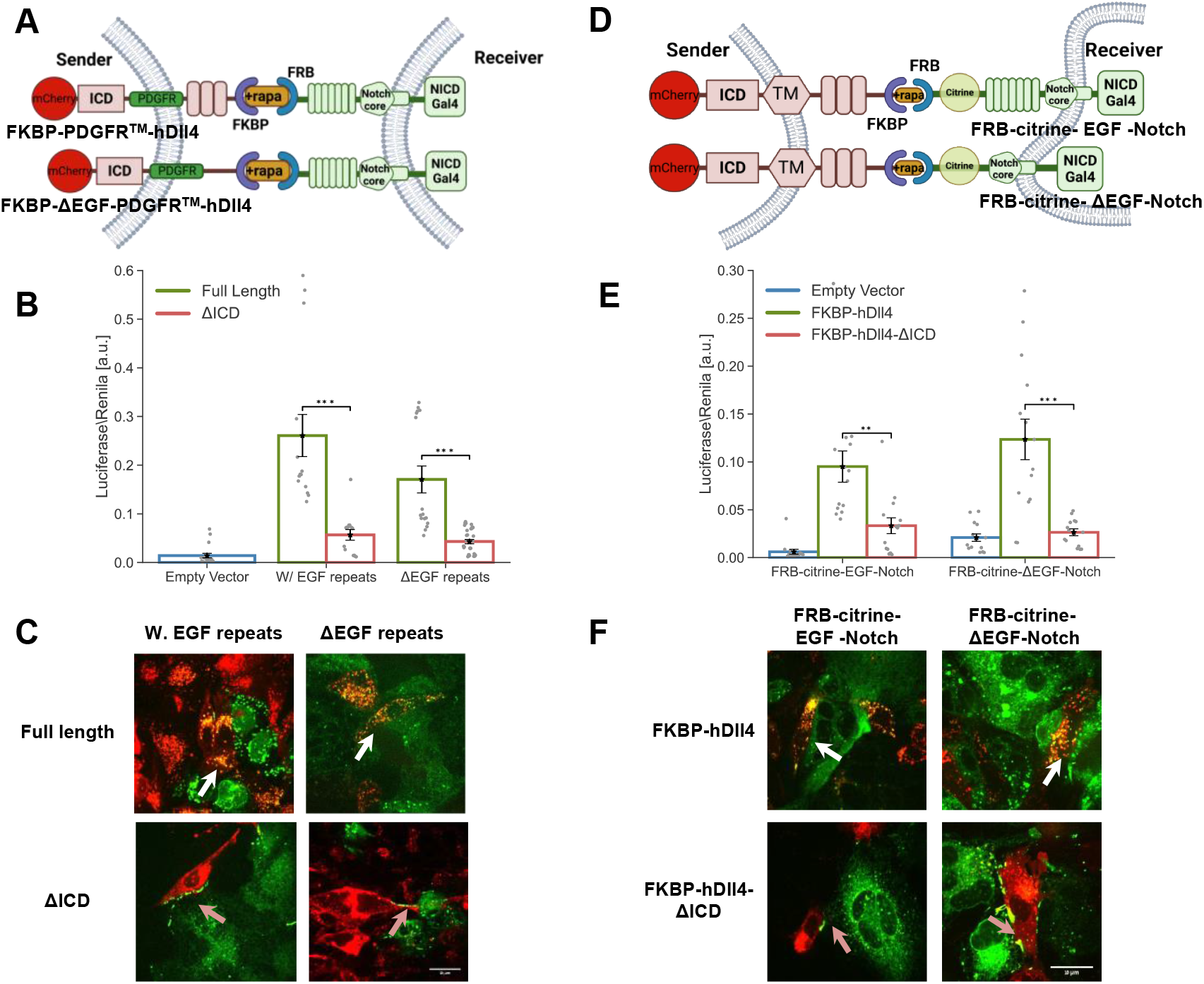
Removing the EGF repeats from the ff-synNotch ligands and receptors does not compensate for the lack of an ICD. **(A, D)** Schematics of the ff-synNotch systems containing either ligands lacking the EGF (1-8) repeats in their ECD (A), or receptors lacking the EGF (24-36) repeats in their ECD (D). **(B, E)** Luciferase activity assay showing the activation of Notch reporter cell lines co-cultured with CHO-TetR cells expressing different ligand variants corresponding to the constructs shown in (A, D). **(C, F)** Images from a TEC assay where U2OS cells expressing Notch-citrine variants were co-cultured with CHO-TetR cells expressing ligand variants corresponding to the constructs shown in (A, D). Images were taken 10 hours induction of ligand expression with 100 ng/ml dox followed by 1 hour induction of binding with 250 nM rapamycin. TEC was observed with full-length ligands (white arrows). Accumulation at the boundaries of cell contact without TEC was observed with ligands lacking their ICD (orange arrows). Data points show mean values from n=14 for (B, E) from 4 independent experiments. Error bars represent S.E.M. ***p<0.001. Scale bars-10μm.

Next, we tested whether removing the EGF repeats from the synNotch receptor can compensate for the lack of ligand ICD (Figure 5D). As with the previous experiment, both luciferase and TEC assays showed that removing the EGF repeats on the receptor side cannot compensate for the lack of ligand ICD (Figures 5E and 5F, and Figure S3). These results show that neither the presence of the EGF repeats in the receptors nor in the ligands can account for the differences between ff-synNotch and aa-synNotch systems.

### Minimal ff-synNotch system requires the ligand ICD

We have shown above that no single structural element (*e*.*g*., TM domain, EGF repeats) can account for the difference between aa-synNotch and ff-synNotch systems. We next tested whether replacing all these domains together to generate a minimal ff-synNotch system could lead to activity without ligand ICD. In these experiments, we used the minimal ff-synNotch receptor lacking the EGF repeats, and we systematically replaced the TM domain and removed the EGF repeats from the ligand ECD (Figures 6A-F), testing these variants in the luciferase and TEC assays. Altogether, the results from these experiments are consistent with the above results showing that the ligand ICD is also required for a minimal ff-synNotch system.

**Figure 6.**
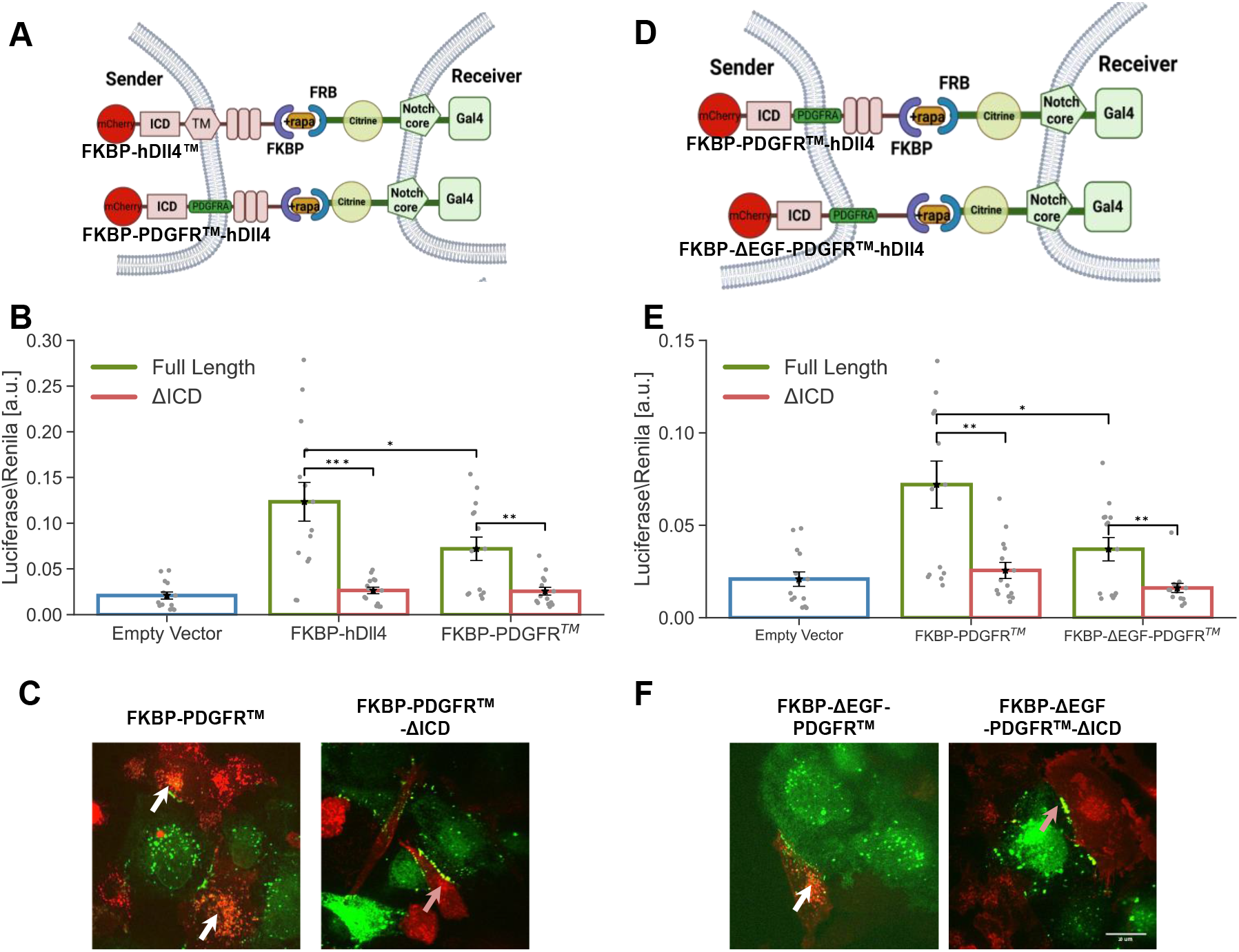
Minimal ff-synNotch systems require the ligand ICD. **(A, D)** Schematics of the minimal ff-synNotch systems where both ligand TM domain (A) and EGF (1-8) repeats (D) are changed as indicated. Receptors lack the EGF (24-36) repeats. **(B, E)** Luciferase activity assay showing the activation of Notch reporter cell lines co-cultured with CHO-TetR cells expressing different ligand variants corresponding to the constructs shown in (A, D). **(C, F)** Images from a TEC assay where U2OS cells expressing Notch-citrine variants were co-cultured with CHO-TetR cells expressing ligand variants corresponding to the constructs shown in (A, D). Images were taken 10 hours induction of ligand expression with 100 ng/ml dox followed by 1 hour induction of binding with 250 nM rapamycin. TEC was observed with full-length ligands (white arrows). Accumulation on the boundaries, but no TEC were observed in ligands lacking their ICD (orange arrows). Data points show mean values from n=14 for (B, E) from 4 independent experiments. Error bars represent S.E.M. *p<0.05,**p<0.01, ***p<0.001. Scale bars-10μm.

## Discussion

The main question investigated in this work is why ligand endocytosis is required for activation in endogenous Notch signaling, but not for activation in the aa-synNotch system. Understanding the origin of this difference is important because it may reveal mechanistic insights into the activation of endogenous Notch on one hand, and concurrently provide guidelines for designing better synNotch systems. We compared the endogenous system to two types of synthetic Notch systems (i.e., the aa- and ff-synNotch systems) and used chimeras and domain swaps to study how different structural properties of synthetic and endogenous ligands and receptors affect their activity. We first confirmed that endogenous Notch does require the ligand ICD for proper activation, while aa-synNotch dos not. Moreover, adding the ligand ICD to the aa-synNotch system did not lead to increased signaling activity. Because ligand ubiquitylation is required to deliver a pulling force to the receptor in the endogenous system (Musse, Meloty-Kapella and Weinmaster, 2012; Gordon *et al*., 2015), it is unclear what replaces that requirement in the aa-synNotch system.

These studies showed that increasing receptor-ligand affinity, changing the identity of the TM, and including or removing EGF repeats from the ECD of the receptors and the ligands cannot compensate for the lack of the ligand ICD in either endogenous or ff-synNotch signaling. This conclusion is buttressed by data from two activity assays, one using a transcriptional reporter and the other using TEC of the receptor ectodomain. Overall, our analysis ruled out that the differences in behavior between aa-synNotch and endogenous/ff-synNotch systems are due to simple structural or molecular properties. These findings suggest that the aa-SynNotch system bypasses the ligand ICD requirement because of a differential ability of antigen-antibody pairs to promote other adhesive cell-cell interactions that provide the mechanical tension needed for ligand activation.

What could then explain the functional differences between the aa-synNotch and endogenous/ff-synNotch? First, it is clear that this difference in behavior is not specific to GFP-αGFP interactions since multiple aa-synNotch systems based on different antibody-antigen pairs have been developed (e.g., CD19-αCD19) (Morsut *et al*., 2016; Choe *et al*., 2021). None of these aa-synNotch systems require the ligand ICD. Second, based on the comparison between minimal ff-synNotch and aa-synNotch, it seems that the difference is not due to the sizes of the extracellular domains of receptor and ligand pairs. However, this does not rule out that different orientations or more complex structures may play a role. While we have not identified here the specific factors that lead to the functional differences, our results significantly narrowed the possible mechanisms that can give rise to these differences. One possible explanation could be attributed to differences in stability or immobilization of receptor-ligand complexes, possibly through cluster formation. It has been suggested that the binding of Notch is more effective when ligands are clustered (Hicks *et al*., 2002), and that recombinant soluble ligands require clustering or immobilization to activate Notch signaling and induce biological responses (D’Souza, Miyamoto and Weinmaster, 2008). In our case, it is possible that the antigen-antibody (i.e., GFP-αGFP) interaction forms clusters more readily than do the endogenous and ff-synNotch systems (Taylor *et al*., 2017).

Finally, it has been demonstrated that mutant ligands lacking the ICD or lysine that are unable to activate Notch undergo reverse TEC (i.e., the ligands go through endocytosis in the receiver cells) (Langridge and Struhl, 2017). Our TEC images indicated a similar behavior in our system when removing the ligand ICD. However, recent work showed that aa-synNotch exhibits reverse TEC in a system that is presumably active (Tang *et al*., 2020). It would be interesting to explore and quantify under what conditions reverse vs. normal TEC occurs and how it affects signaling activity.

It is already known that Notch ligands show significant diversity in their affinity for Notch receptors. For example, Dll4 binds to the Notch1 with higher affinity than either Dll1(Andrawes *et al*., 2013) or Jag1 (Luca *et al*., 2015), in a manner that is modulated by the expression of the glycosyl transferases from the Fringe family (Takeuchi *et al*., 2018). Hence, receptor-ligand affinity is itself modulated by post-translational modifications that can regulate the ligand activity and potentially control it in different contexts. Our quantitative analysis shows that higher affinity ligands indeed lead to higher activation. However, the higher affinity does not compensate for the lack of ICD (i.e., we observed no activation when the ICD was removed). It will be interesting to determine whether the same relationship is maintained for other Notch receptors (i.e., Notch2-4).

An important application for SynNotch systems is the design of tailored functions in T-cell engineering. Our current research suggests that the mode of activation for aa-synNotch may be different than that of endogenous Notch, because it does not require ubiquitylation of the ligand in the sender cell. It will be important in the future to understand better this mode of activation in order to carry out rational engineering of highly sensitive and inducible next generation synNotch systems.

## Acknowledgments

B.K. is grateful to The Edmond de Rothschild Foundation (IL) for funding her Ph.D. scholarship.

## Author Contributions

This study was conceived and planned by B.K, S.C.B., and D.S. Experiments were performed and analyzed by B.K. Reagents and advice were provided by V.C.L. The manuscript was written by B.K, S.C.B., V.C.L. and D.S.

## Methods

### Cells, plasmids, and reagents

The cell lines used for this study are (1) Chinese Hamster ovary cells (CHO-K1, ATCC-CCL-61) integrated with TetR (Life Technologies, CHO-TetR). Organism: Cricetulus griseus. Sex: female. (2) Human embryonic kidney 293T cells (HEK293T, ATCC CRL3216). Organism: Homo sapiens, human. Sex: Female. (3) Human Bone Osteosarcoma Epithelial Cells (U2OS Line) were kindly provided by Stephen C. Blacklow (Harvard Medical School).

CHO cells were grown in Minimum Essential Medium Eagle - alpha modification (αMEM), HEK293, and U2OS cells were grown in adherent cultures in Dulbecco’s Modified Eagle’s Medium (DMEM), the medium supplemented with 10% FBS. All cells were cultured in a humidified atmosphere of 5% CO_2_ at 37°C. Stable and transient transfections were performed using *Trans*IT-LT1 reagent (Mirus Bio) or Lipofectamine 3000 (Thermo Fisher Scientific) according to the manufacturer’s instructions.

For the Dll constructs, 300ng of the plasmid was taken with an empty vector (800 ng). In brief, cells were transfected with Dll construct, after two days transferred to a 6-well plate, and placed under selection for 100ng/ul Hygromycin or 400ug/ml Zeocin (Invivogen) for two weeks. Single colonies are then isolated using limiting dilution. Colonies were picked up and tested for fluorescence and activity. To reduce clonal variability, we generated a mixed population containing several single clones (typically 3-8 single clone colonies).

All genetic constructs were constructed using standard cloning techniques. Tagged hDll1/4 variants and Notch1-citrine (Figures 1a-d) are based on constructs developed in (Khait *et al*., 2016; Shaya *et al*., 2017). In Notch1 variants, the citrine tag was inserted prior to the NRR domain (between G1435 and A1436). The GFP-αGFP aa-synNotch AAV constructs (Figures 1E-F) are based on (Morsut *et al*., 2016) hDll4ICD was added to the c-terminus of the GFP ligand. The high affinity Dll4 chimeras (Figure 2) were constructed by fusing the TM and ICD of hDll4 to the ECD of the high affinity ligands developed by (Luca *et al*., 2015). The ff-synNotch constructs (Figure 3) were developed based on (Gordon *et al*., 2015). A mCherry was added to the c-terminus of the FKBP ligand, and citrine (for the TEC experiments) was added between the FRB domain and the EGF repeats. Constructs containing the variants with PDGFRb TM domain (Figures 4-6) were constructed from either hDll4 or FKBP ligands. The Dll4 TM was replaced by the PDGFRb sequence from the GFP ligand of the aa-synNotch. All plasmids used in this study are listed in Supplementary table 1 (also contains links to full sequences).

### Lentivirus Production

We used lentivirus to generate U2OS stable cell lines expressing either GFP ligand, GFP-ICD ligands, αGFP receptor, FRB-Citrine-Notch1-Gal4, and FRB-Citrine-ΔEGF-Notch1-Gal4. Lentivirus was produced by co-transfecting the PHR plasmids and vectors encoding packaging proteins (pMD2.G and PsPax2) using the LT1-transfection reagent in HEK293T cells plated in 6-well plates at approximately 70% confluence. Viral supernatants were collected 2 days after transfection and 0.45 μm filtered. The supernatant was used for transduction immediately.

### Luciferase Activity Assay

The activity of the different ligands was tested using a luciferase reporter gene assay. Receiver cells stably expressing either Notch1 or synNotch variants were co-transfected in 24-well plate using TransIT-LT1 (Mirus) or Lipofectamine 3000 (Thermo Fisher Scientific, for U2OS cells) with a Gal4-firefly luciferase reporter (Andrawes et al., 2013) (300 ng) and pRL-SV40 Renilla luciferase (10 ng). 24 hours after transfection, the cells were trypsinized and co-culture with cells stably expressing the ligands. 48 hours after plating, firefly luciferase and Renilla luciferase activities were measured by luminometer (Veritas). Cells were lysed with 100 μl/well Passive lysis buffer x1 (Promega) for 10 min. 20μl of each sample was used for luciferase activity using filtered luciferase buffer including: 26 mg of (MgCO3)4 Mg(OH)2 (Sigma), 20 mM Tricine (Sigma), 0.1 mM EDTA (Biological Industries), 2.67 mM pH=7.8 MgSO4 (Merck). For the luciferase reaction, we used luciferase buffer supplemented with 0.4 mM ATP (Sigma), 26.6 mM DTT (Sigma), Coenzyme A X0.8 (Sigma), and 0.4 mM D-Luciferin, and for Renilla activity using filtered Renilla buffer including 80 mM di-Potassium hydrogen phosphate trihydrate (Merck) and 20 mM Potassium dihydrogen phosphate for analysis (Merck). Notch activity is expressed as a ratio of normalized luciferase by Renilla.

### TEC assay

For the TEC assay, sender and receiver cells with a 1:1 ratio were seeded 24 hours before imaging in 24-well glass bottom plates (De-Groot). Directly prior to imaging, the media was replaced with low fluorescence imaging media (αMEM without Phenol red, ribonucleosides, deoxyribonucleosides, folic acid, biotin, and vitamin B12 (Biological Industries, Israel) and 100ng/ml doxycycline (Sigma-Aldrich) was added to the growth medium to induce ligand expression. For the ff-synNotch system, 250nM/ml Rapamycin (Zotal) was added as well.

### Microscopy Details

Cells were imaged using Andor revolution spinning disk confocal microscope with DPSS CW 515nm and 561nm and 488nm 50mW lasers (Andor, Belfast, Northern Ireland). The imaging setup consisted of an Olympus inverted microscope with an oil-immersion Plan-Apochromatic 60x objective NA=1.42 (Olympus, Tokyo, Japan); and an ANDOR iXon Ultra EMCCD camera (Andor, Belfast, Northern Ireland). The microscope was equipped with a 37 °C temperature-controlled chamber and a CO_2_ regulator providing 5% CO_2_ (Okolab, Italy). The equipment was controlled by Andor iQ software (Andor, Belfast, Northern Ireland).

### Flow Cytometry

CHO and U2OS cells were seeded in 6-well plates at approximately 70% confluence 24 hours before FACS. Cells were treated with 100ng/ml doxycycline right after seeding. Directly prior to FACS, cells were trypsinized, spun at 1000 rpm for 5 min, and resuspended in 200 μL of FACS buffer. FACS buffer consisted of PBS with 1% FBS serum and 5mM EDTA. Flow cytometry was performed using a Cytoflex5L flow cytometer (Beckman Coulter). Kaluza software was used to analyze the data.

## Supplemental information

**Figure S1.**
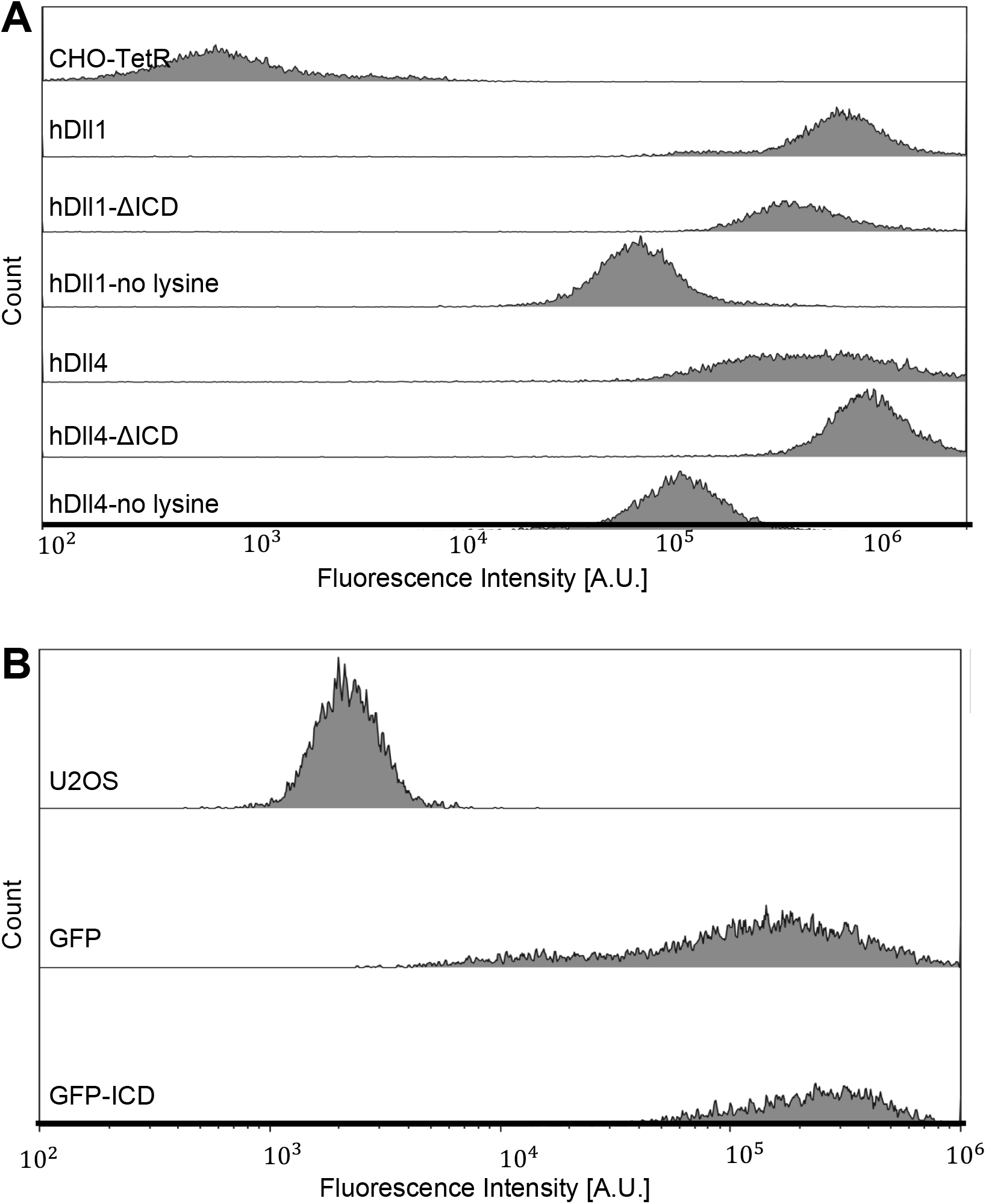
FACS analysis of fluorescently tagged ligands. **(A)** Comparison of the fluorescence of CHO-TetR cells expressing different ligand variants tagged with mCherry (as indicated for each row). Control cells (CHO-TetR) that do not express fluorescent protein are shown in the top row. **(B)** Comparison of the fluorescence of U2OS cells expressing either the GFP or GFP-ICD ligand of the aa-synNotch system (as indicated). Control cells (U2OS) that do not express fluorescent protein are shown in the top row.

**Figure S2.**
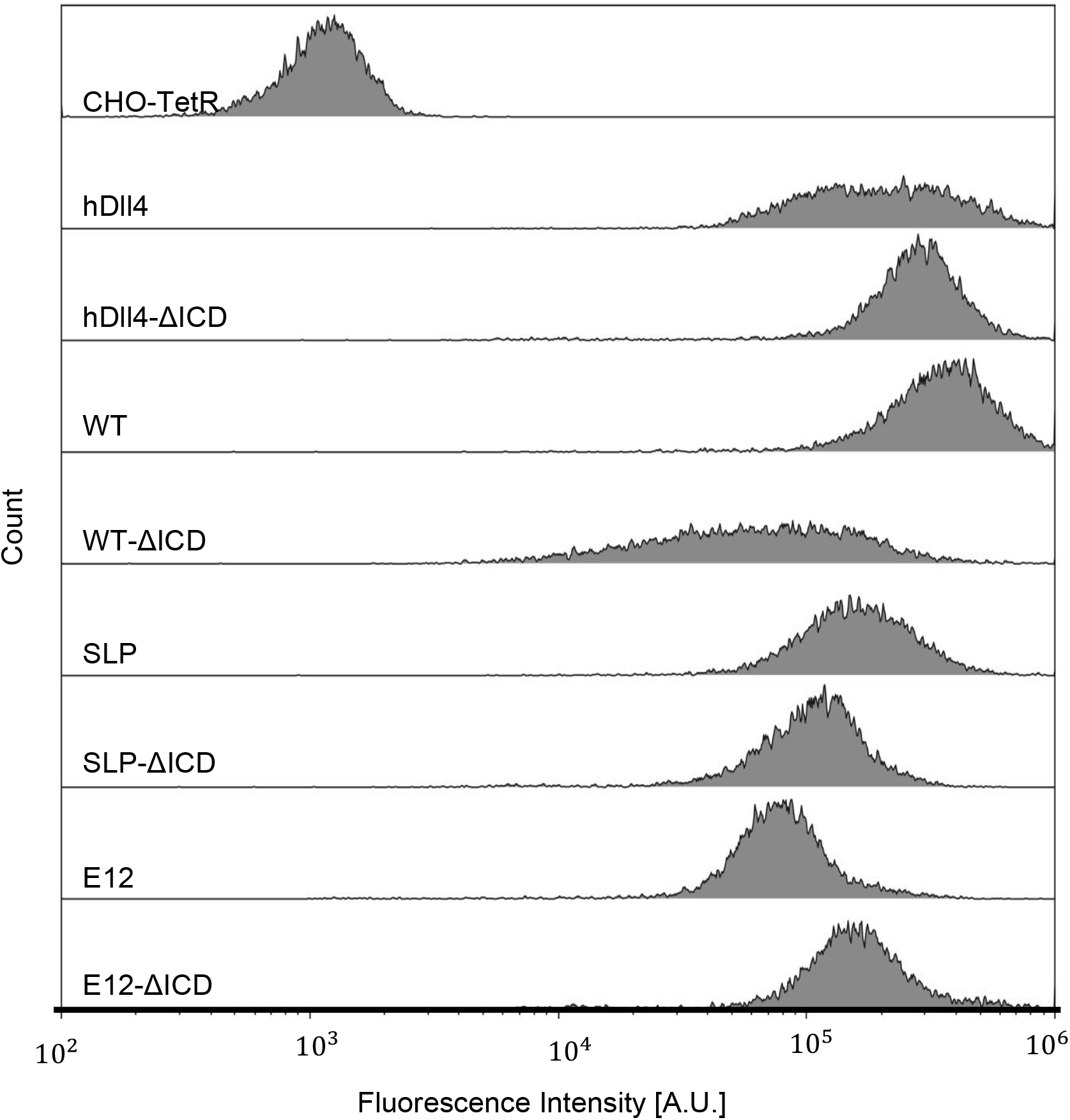
FACS analysis of fluorescently tagged ligands. Comparison of the fluorescence of CHO-TetR cells expressing different ligand variants tagged with mCherry (as indicated for each row). Control cells (CHO-TetR) that do not express fluorescent protein are shown in the top row.

**Figure S3.**
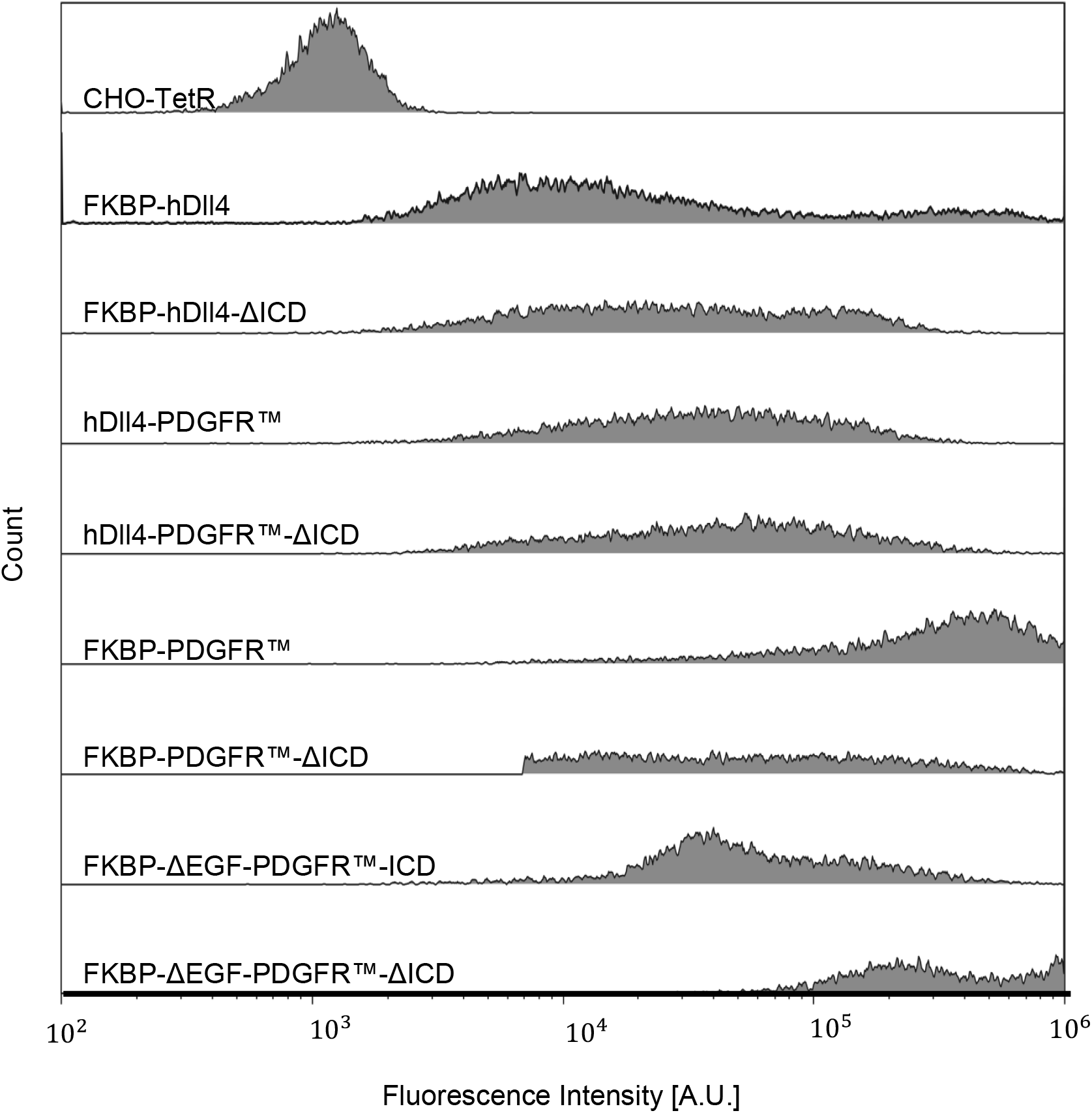
FACS analysis of fluorescently tagged ligands. Comparison of the fluorescence of CHO-TetR cells expressing different ligand variants tagged with mCherry (as indicated for each row). Control cells (CHO-TetR) that do not express fluorescent protein are shown in the top row.

